# Understanding the Role of Pyruvate Dehydrogenase in *Listeria monocytogenes* Virulence

**DOI:** 10.1101/2025.05.29.656449

**Authors:** Matthew J. Freeman, Noah J. Eral, Abigail M. Debrine, David M. Stevenson, Daniel Amador-Noguez, John-Demian Sauer

## Abstract

To survive within restrictive host niches, bacterial pathogens must possess finely tuned physiological adaptations. One such niche inhabited by *Listeria monocytogenes* (*L. monocytogenes*) is the host cell cytosol—a compartment characterized by significant barriers to entry, metabolic limitation, and immune surveillance. Previously, we identified *L. monocytogenes* transposon mutants defective for intracellular survival due to disruptions in key metabolic pathways, including cell wall biosynthesis, menaquinone production, and pyruvate metabolism. One of these mutants mapped to a central component of the pyruvate dehydrogenase (PDH) complex, *pdhC*::Tn. Notably, this mutant exhibits pronounced survival defects during infection, despite retaining robust growth and survival in nutrient-rich media. We go on to show that disruption of *pdhA*::Tn and *pdhD*::Tn similarly led to virulence attenuation during intra-macrophage growth, plaquing assays, and murine infections. Respiro-fermentative metabolic profiling revealed that *pdhC*::Tn mutants have an altered respiro-fermentative metabolism with more prominent secretion of lactate. Further, unbiased metabolomic profiling revealed a global starvation phenotype with lower levels of upper glycolytic intermediates and TCA cycle intermediates coupled with elevated intra-bacterial levels of pyruvate and lactate. We then demonstrate that PDH mutants are unable to efficiently utilize phosphotransferase (PTS)-dependent carbon sources and that their growth can be rescued using non-PTS-mediated carbon sources such as hexose phosphates. To identify genetic suppressors of PDH deficiency, we performed an EMS mutagenesis screen using fructose—a PTS-transported carbon source—as the sole carbon source. Five suppressors each contained a single independent mutation in the redox sensing regulator *rex*. Subsequently, we show that loss of *rex* restores *pdhC*::Tn’s ability to consume PTS-mediated carbon sources through the alleviation of fermentative repression. Further, *pdhC*::Tn suppressor mutants show restored intracellular growth, but not virulence *in vivo*. Together, these findings indicate that a key defect in PDH mutants is the inability to import and metabolize PTS-dependent carbon sources in the host cytosol. We posit this impairment leads to disruptions in redox balance and a shift in respiro-fermentative metabolism, ultimately contributing to the loss of intracellular fitness and virulence.

## INTRODUCTION

*Listeria monocytogenes* (*L.* monocytogenes) is a Gram-positive, cytosolic bacterial pathogen capable of causing severe morbidity and mortality (1–3). It is well established that for *L. monocytogenes* to successfully infect and cause disease, it must invade host cells, and survive in the restrictive host cytosol (4–8). To access the cytosol, *L. monocytogenes* employs a well-characterized arsenal of virulence factors under the control of its master regulator, PrfA (9,10). Key among these is listeriolysin O (LLO), a pore-forming toxin that targets cholesterol-containing membranes and enables escape from the acidified, toxified phagolysosome (11–13). Once in the cytosol, *L. monocytogenes* induces expression of its hexose phosphate transporter, UhpT, and hijacks host actin polymerization machinery via ActA to facilitate intracellular movement and spread to neighboring cells (14–16). While the canonical virulence factors supporting the intracellular lifecycle of *L. monocytogenes* are well defined, much less is known about the metabolic genes and pathways that support survival in this unique niche (17). More specifically, it is vital to understand how *L. monocytogenes* has adapted its metabolism to the cytosol as this can inform barriers to bacterial survival in the cytosol and potential antimicrobial targets (4,18,19).

To investigate genes essential for *L. monocytogenes’* cytosolic survival, our lab previously executed a forward genetic screen to identify *L. monocytogenes* mutants that are killed in the macrophage cytosol (20). We identified mutants with defects in cell wall biosynthesis, menaquinone synthesis, and pyruvate metabolism (20). Interestingly, these mutants showed no growth defects in rich media, suggesting that their attenuation results from specific vulnerabilities to host cytosolic conditions rather than general physiological impairment. Why pyruvate metabolism, specifically the pyruvate dehydrogenase (PDH) complex, is required for cytosolic survival but not *in vitro* viability, remains unresolved (21–23).

In *L. monocytogenes*, the PDH complex consists of four subunits, encoded in a single operon, that form a large multiprotein complexes that converts pyruvate into acetyl CoA. (24). The E1 subunit is composed of two proteins encoded by *pdhA* (LMRG_00514) and *pdhB* (LMRG_00515), which decarboxylate pyruvate to form an active acetaldehyde intermediate bound to thiamine pyrophosphate (TPP). The E2 subunit, encoded by *pdhC* (LMRG_00516), then catalyzes transacetylation to CoA and reduces lipoic acid. The E3 subunit, encoded by *pdhD* (LMRG_00517), reoxidizes lipoic acid and transfers electrons to NAD⁺, generating NADH. In sum, PDH irreversibly converts pyruvate to acetyl-CoA during aerobic metabolism through tightly coordinated steps that minimize dilution of intermediates and off-target reactions (25). Additionally, PDH enzymatic activity is allosteric regulated; it is stimulated by phosphoenolpyruvate (PEP) and AMP, and inhibited by NADH and acetyl-CoA, presumably to define when energy sources are low, but resources are high (25,26). Because PDH sits at the metabolic junction between glycolysis and the tricarboxylic acid (TCA) cycle, its disruption is likely to result in broad, pleiotropic effects. For example, PDH mutants are deficient in acetyl-CoA production, a precursor essential for fatty acid biosynthesis (27). In addition, reduced TCA cycle flux may impair NADH generation necessary for electron transport chain function and the synthesis of TCA-derived amino acids (14,28). Thus, analysis of PDH-deficient mutants and their virulence phenotypes must consider this range of interconnected metabolic disruptions.

In evaluating these pleiotropic defects, it is also critical to assess the carbon sources being acquired and funneled into the PDH complex, as well as its broader impact on cellular redox and energy states. Our lab recently demonstrated that intracellular *L. monocytogenes* is only modestly reliant on glycerol and hexose phosphates, and instead depends heavily on phosphotransferase systems (PTS) to acquire host-derived carbon sources (29). Importantly, this PTS function is tightly linked to pyruvate metabolism, as it terminally relies on the conversion of PEP to pyruvate to transfer phosphate groups via EI (*ptsI*) and HPr (*ptsH*) to substrate-specific EII complexes (25,30–32). These transporters phosphorylate incoming sugars, which then enter upper glycolysis (25). Interestingly, PTS-encoding genes are enriched in bacteria that are facultative or strict anaerobes and PTS appear to be less common among strict aerobes (33,34). It has been hypothesized this evolutionary selection is because for PTS to import and phosphorylate a new carbohydrate, a molecule of PEP is used; thus, leaving only one PEP from glycolysis for biosynthetic pathways (33,35). In aerobic organisms, this PEP will be rapidly catabolized, while anaerobic organism can more readily retain PEP for essential biosynthetic purposes (33,34). Some evidence in the field suggest the *L. monocytogenes* splits its carbon use of different metabolites for either anabolic or catabolic processes (36,37). However, recent evidence suggests these previously described carbon sources are not essential for infection and therefore the relevance of this model remains undetermined (29). Thus, for simplicity, from initial phosphorylation, carbon is funneled through glycolysis and into the TCA cycle, generating ATP, NADH, and FADH₂ (25). Under aerobic conditions, these reduced cofactors support oxidative phosphorylation and ATP synthesis. Importantly, cells have evolved highly sophisticated methods of detecting the successful balance of NADH and NAD+ (23,38), a process particularly important for bacteria that have both the ability to ferment and respire as they must be able to modulate between these states to promote the most efficient use of carbon and deal with unique environmental stressors (23,39,40).

Bacterial pathogens, including *L. monocytogenes*, employ metabolic strategies that allow them to utilize carbon while evading host cell pressures and immune detection (41–43). One example of host defenses targeting bacterial metabolism is the use of reactive nitrogen species (RNS) by macrophages, which inhibit bacterial aerobic respiration (44,45). From the pathogen side, some bacterial species—including *Salmonella enterica* and *Staphylococcus aureus*— require aerobic respiration for full virulence (46–50). *L. monocytogenes*, on the other hand, is known to dynamically modulate its metabolism between fermentation and respiration, often using both simultaneously in what is referred to as respiro-fermentative metabolism (23). During respiration, *L. monocytogenes* regenerates NAD⁺ from NADH by unloading electrons into the electron transport chain (ETC) (23). For which *L. monocytogenes* employs two respiratory chains: one utilizing oxygen as the terminal electron acceptor, and another using extracellular ferric iron and fumarate (22,23,51,52). Under aerobic conditions, *L. monocytogenes* shifts its fermentative output from lactate to acetate in order to maximize ATP production, albeit at the expense of NADH regeneration (53). To balance this redox requirement, a portion of carbon cannot be fully oxidated and continues to be fermented to lactate, which supports NADH regeneration but yields less ATP. This metabolic modulation results in markedly different carbon consumption profiles and distinct metabolic by-products, which can be used to infer the bacterium’s metabolic state: elevated lactate levels indicate purely fermentative metabolism, while acetate production is associated more with respiro-fermentative state (23,53).

To solve the challenge of balancing metabolic demand and redox homeostasis, bacterial pathogens have acquired sophisticated muli-layered method of sensing and adjusting their metabolism to respond to these demands (38). Some of this sensing occurs by enzymes that require these cofactors such as dehydrogenase complexes (25,26). One advantage of this model is it allows enzyme to be at the ready for their respective functions while detecting the metabolic state of the cell. One downside is that it is highly costly for bacteria to produce and retain these enzymes if they are not functional. To overcome this limitation bacteria also utilize regulators and metabolic sensors to control the production of the enzymes and thus control metabolism. *L. monocytogenes* and other pathogens encode Rex, a redox-sensing transcriptional regulator that responds to intra-bacterial NAD⁺/NADH ratios (38,54,55). When NAD⁺ levels are low, Rex derepresses genes involved in fermentation (54). Conversely, high NAD⁺/NADH ratios lead to repression of key fermentation genes. While the full significance of Rex-mediated regulation remains unclear, preliminary data suggest that *rex* mutants of *L. monocytogenes* are modestly attenuated during oral infection in murine models but retain normal *ex vivo* growth within macrophages (38).

In this study, we evaluate the contribution of pyruvate dehydrogenase (PDH) deficiency to *L. monocytogenes* virulence. We find that mutants lacking any individual PDH component (*pdhA*, *pdhC*, or *pdhD*) exhibit equivalent defects in virulence, indicating that loss of any one component completely ablates the function of the complex. *pdhC::Tn* mutants show an altered respiro-fermentative metabolism with a shift from acetate production toward that of lactate.

Metabolomic profiling of a *pdhC::Tn* mutant revealed global depletion of upper glycolytic and TCA cycle intermediates, accompanied by an accumulation of pyruvate and lactate when grown in rich media. We further demonstrated that PDH mutants are defective in utilizing phosphotransferase system (PTS)-transported carbon sources. However, their growth could be restored in defined media supplemented with hexose phosphates (+glutathione). A suppressor screen identified five independent suppressor mutations in the gene encoding the redox-sensing transcriptional regulator, Rex. All of these suppressor mutations restored *pdhC::Tn* mutant growth on the PTS-dependent carbon source fructose. One suppressor, which introduced a premature stop codon in *rex*, restored growth of *pdhC::Tn* mutants *ex vivo* in macrophages, though it did not rescue virulence *in vivo*. Collectively, our findings demonstrate that *L. monocytogenes* requires rapid conversion of pyruvate to acetyl-CoA via PDH to sustain both redox homeostasis and PTS-dependent carbon acquisition—two processes essential for intracellular growth and pathogenesis. At least part of the virulence defect in macrophages for PDH mutants is attributable to an inability to utilize PTS-transported carbon sources, likely due to Rex-mediated repression of fermentation under redox-imbalanced conditions.

## RESULTS

### Pyruvate dehydrogenase mutants are significantly attenuated for intracellular growth and virulence, while maintaining WT levels of growth in rich media

The *pdhC::Tn* mutant identified in our previous work exhibits significant virulence defects across multiple assays (20). Notably, this mutant was completely unable to grow within the cytosol of bone marrow-derived macrophages and was actively cleared, as indicated by decreasing bacterial burdens over time and bacteriolysis in the macrophage cytosol (20). During acute murine infection, the *pdhC::Tn* mutant was fully attenuated, with bacterial burdens falling to the limit of detection by 48 hours post-infection (20). In contrast, L2 fibroblast plaquing assays—which assess both intracellular growth and cell-to-cell spread—revealed that the *pdhC::Tn* mutant retained partial function, forming plaques approximately 50–70% the size of those formed by wild-type *L. monocytogenes* (20). These findings suggest that the pyruvate dehydrogenase complex (PDH) is critical for intracellular survival and full virulence of *L. monocytogenes*, however, it remained unclear whether these phenotypes were specific to the E2 subunit (PdhC) or were generalizable to other subunits of the PDH complex.

First, we hypothesized that loss of any individual component of the PDH complex would result in similar virulence defects, but like *pdhC*::Tn would retain the ability to grow in rich media. To test this, we obtained unpublished transposon mutants in *pdhA* and *pdhD* from Dr. Daniel Portnoy (University of California, Berkeley) and conducted a series of standard *in vitro* growth curves and virulence assays to determine the contributions of individual PDH subunits. Notably, a *pdhB* mutant has never been isolated from bacterial forward genetic screens, perhaps due to functional redundancy with the E1β subunit of the branched chain keto-acid dehydrogenase complex which shares 62% identity and ∼75-80% similarity with the PDH E1β subunit (56,57). *In vitro* growth curves of PDH mutants in rich media demonstrated that *pdhA*::Tn, *pdhC*::Tn and *pdhD*::Tn mutants readily grow in rich media (**Figure 1A**), again highlighting that PDH mutant defects during virulence are specific to *L. monocytogenes* physiology during infection. Next, we performed intracellular growth curves in bone marrow-derived macrophages (BMDMs), cells that *L. monocytogenes* uses as it primary *in vivo* niche, to assess cytosolic invasion and replication over an 8-hour period (58,59). All available PDH subunit mutants were unable to grow and were cleared from the host cytosol, mirroring the phenotype of the original *pdhC::Tn* mutant (**Figure 1B**). This supported that loss of any one component of the PDH complex would result in the inability to survive and replicate in the cytosol.

**Figure 1.**
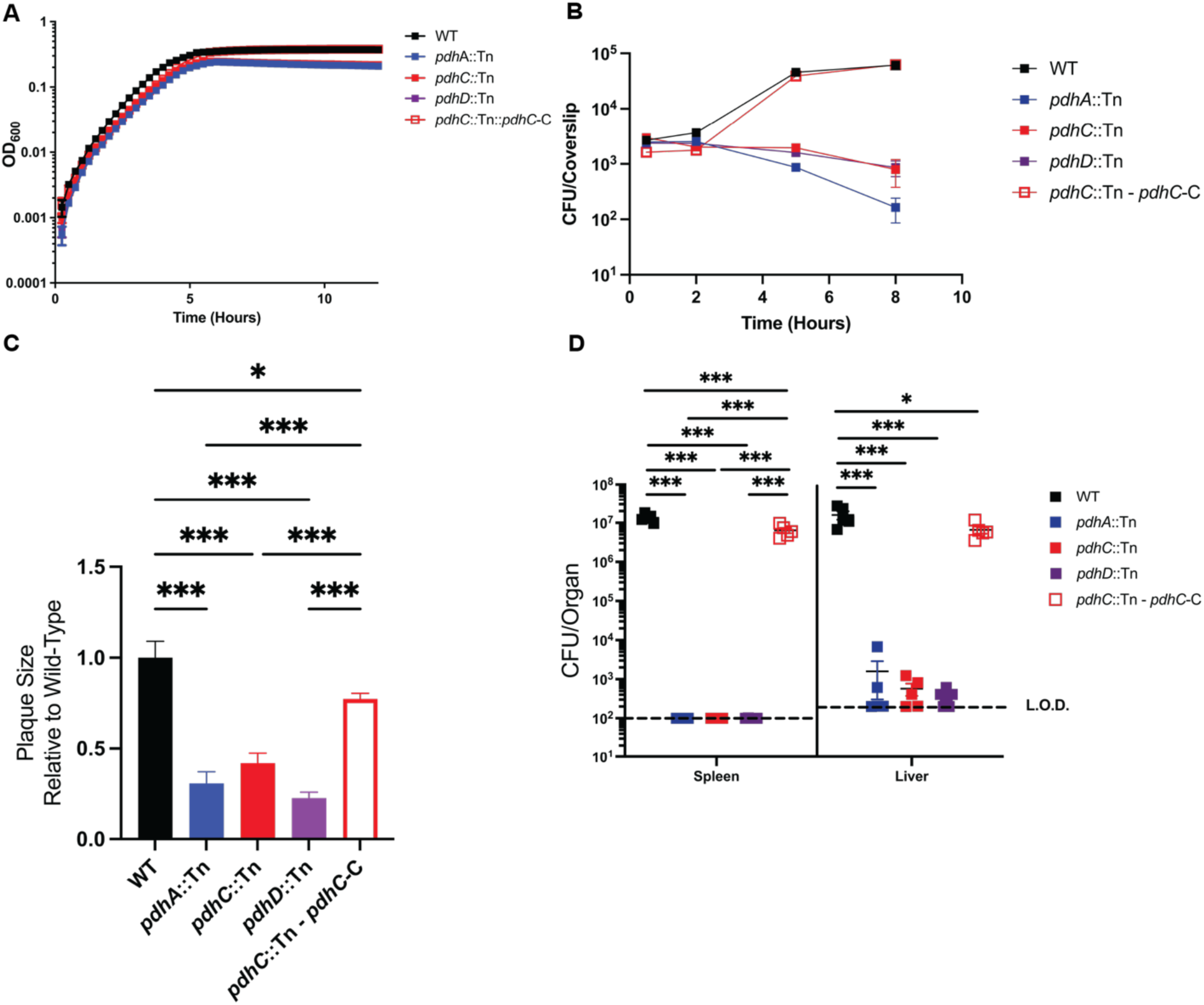
PDH complex mutants retain wild-type growth in rich media and have phenotypically similar virulence defects. (A) Indicated strains were grown in Brain, Heart Infusion (BHI) at 37°C, shaking at 250 r.p.m. and had OD600 measured every 15 minutes for 12 hours in a plate reader. (B) Intracellular growth of indicated strains was determined in BMDMs following infection at an MOI of 0.2. Growth curves are representative of 3 independent experiments. Error bars represent the standard error of the means of technical triplicates within the representative experiment. (C) L2 fibroblasts were infected with indicated *L. monocytogenes* strains at an MOI of 0.5 and were examined for plaque formation 4 days post infection. Assays were performed in biological triplicate and data displayed is the Mean and SEM of a strain’s plaque size relative to WT in one of three representative biological replicates. (D) Bacterial burdens from the spleen and liver were enumerated at 48 hours post-intravenous infection with 1×10^5^ bacteria. Data are representative of results from two experiments. Horizontal dashed lines represent the limits of detection, and the bars associated with the individual strains represents the mean and SEM of the group.

While intramacrophage growth curves measure invasion, single cycle infection and growth, plaque assays offer insight into bacterial virulence across prolonged periods of growth and the ability to spread to neighboring cells. Therefore, we evaluated the requirement of each PDH subunit for *L. monocytogenes* virulence during L2 fibroblast plaquing assays, predicting that *pdhA* and *pdhD* mutants would retain some ability to grow and spread. While all mutants of the PDH complex were attenuated relative to WT *L. monocytogenes,* there was no statistically significant difference between individual subunit mutants (**Figure 1C**). Together this data shows that PDH mutants share virulence phenotypes across assays, but do perhaps have minorly different virulence capabilities during more prolonged multi-cycle infections versus single cycle infections in macrophages.

Ultimately, to assess virulence *in vivo* under physiologically relevant metabolic and immune pressures, we performed acute murine infection and organ burden assays. In short, C57BL/6 mice were infected intravenously with 1 × 10⁵ CFU of each strain, and spleens and livers were harvested 48 hours post-infection to quantify bacterial burdens. All PDH complex mutants displayed severe attenuation, with bacterial counts near or below the assay’s detection limit in the spleens. Notably, each of the PDH complex mutants displayed occasional very low levels of bacterial burdens in the livers, displaying an organ specific sensitivity (**Figure 1D**).

In summary, our results demonstrate that loss of any PDH complex subunit leads to comparable virulence defects, suggesting a shared physiological basis (**Figure 1A-D**). Further, all defects of *pdhC*::Tn could be rescued to near WT levels with heterologous overexpression of *pdhC* (**Figure 1A-D**). Moving forward we sought to investigate the mechanism of *L. monocytogenes’* virulence attenuation from PDH complex deficiency using *pdhC*::Tn as a representative mutant.

### *PdhC*::Tn mutants show altered respiro-fermentative metabolic byproduct secretion relative to that of WT *L. monocytogenes*

Due to an incomplete tricarboxylic acid (TCA) cycle, *L. monocytogenes* relies on a respiro-fermentative metabolism in which pyruvate is predominantly directed toward fermentative acetate production (53). Previously, our lab identified that *Listeria monocytogenes* mutants deficient in menaquinone biosynthesis—and therefore lacking functional respiratory chains—exhibit substantial alterations in their respiro-fermentative metabolism marked by increased lactate production relative to acetate, indicating a shift toward fermentative metabolism over oxidative metabolism (20–23,60). Further this fermentative byproduct shift has also been observed in *aro* mutants similarly defective for respiration (60). We hypothesized that PDH complex mutants may display similar metabolic alterations due to their impaired ability to efficiently channel carbon from glycolysis into the tricarboxylic acid (TCA) cycle resulting in insufficient levels of NADH for the electron transport chain. To test whether the inability to funnel carbon into the TCA resulted in an altered respiro-fermentative metabolism, we grew each strain overnight in rich media (brain, heart infusion) and analyzed the bacterial supernatants and standards via high-performance liquid chromatography (HPLC) to quantify the relative and absolute abundance of acetate and lactate.

As expected, WT *L. monocytogenes* showed a strong predominance toward the production of acetate versus lactate (**Figure 2**). HPLC of *pdhC*::Tn supernatants revealed that, like other respiration-deficient strains, PDH mutants produce significantly altered respiro-fermentative profiles (**Figure 2 and Supplemental Figure 1B**) (21–23,60). Specifically, PDH mutants exhibited a marked increase in lactate production with a corresponding decrease in acetate, suggesting a shift away from oxidative metabolism with acetate fermentative byproduct production (**Figure 2**). This phenotype could be restored to that of WT *L. monocytogenes* through the heterologous overexpression of *pdhC* (**Figure 2**). Taken together this data demonstrates that loss of PDH results in a disruption in the respiro-fermentative metabolism of *L. monocytogenes in vitro,* marked by a shift to lactate production.

**Figure 2.**
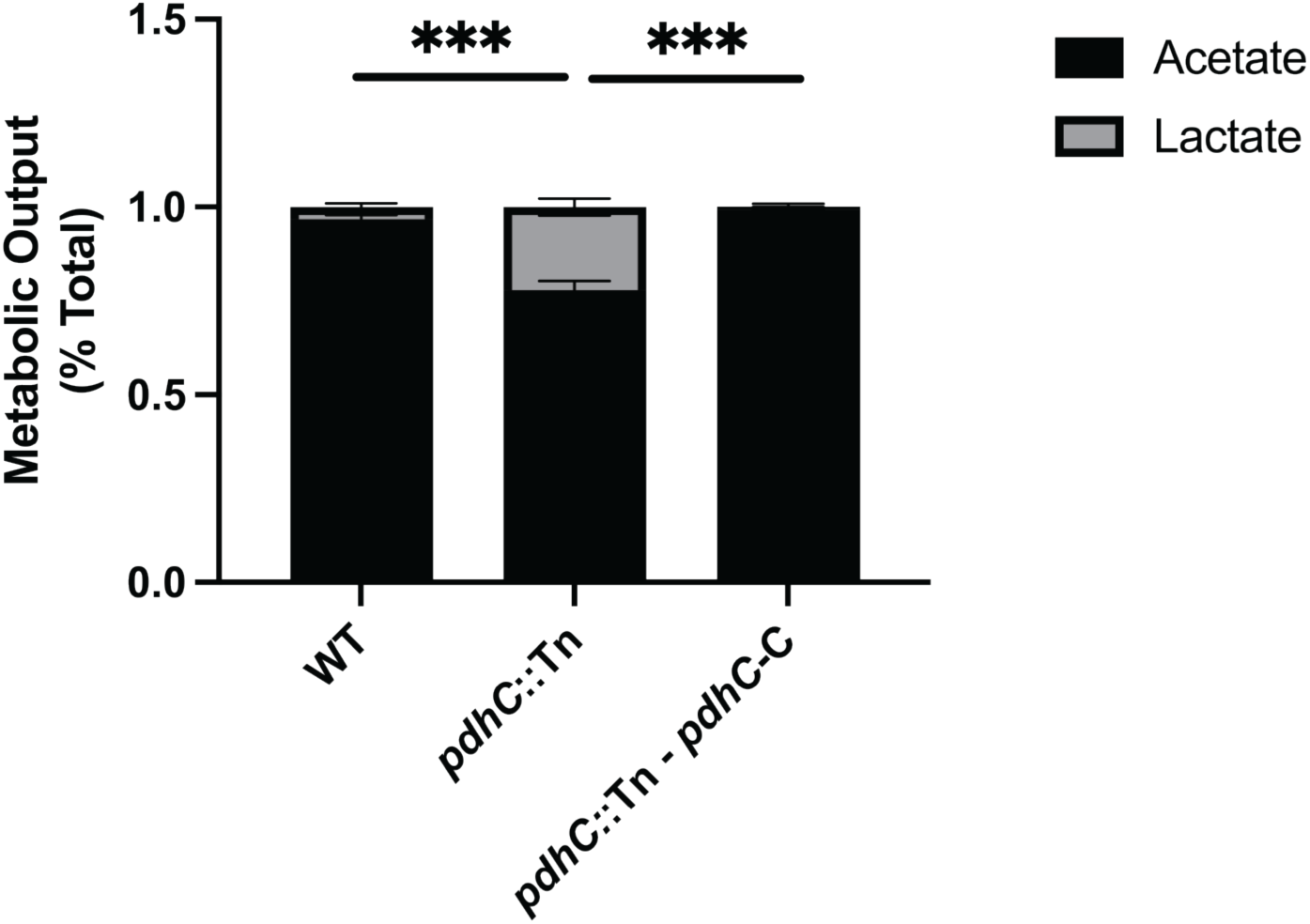
*PdhC*::Tn mutants show altered respiro-fermentative metabolite byproducts relative to WT *L. monocytogenes*. High-performance liquid chromatography (HPLC) was used to quantify fermentation products (acetate and lactate) produced and secreted by the indicated *L. monocytogenes* strains grown aerobically in BHI medium at 37°C to stationary phase. The mean percentage of acetate and lactate production by each strain was compared to that of the wild-type *L. monocytogenes*.

### Pyruvate dehydrogenase mutants are not rescued by restoration of NAD+ production using NOX

We previously demonstrated that one of the primary defects in *L. monocytogenes* menaquinone mutants that contributes to a loss of virulence is an impaired ability to regenerate NAD+ (23). This deficiency is largely attributed to their inability to oxidize NADH via electron transfer through the respiratory chain resulting in a shift in fermentative metabolism toward lactate. In menaquinone-deficient strains, this redox imbalance can be rescued through overexpression of the water-forming NADH oxidase (NOX), which facilitates NADH oxidation independently of the respiratory chain (23). Importantly, NOX overexpression restores virulence in menaquinone mutants across multiple assays, including intracellular growth curves, fibroblast plaquing, and murine infection models (23).

Based on these prior findings and the observation that *pdhC*::Tn mutants show a respiro-fermenatative metabolism shifted toward lactate, we hypothesized that *pdhC*::Tn mutants suffer from NAD⁺ depletion due to impaired flux through the tricarboxylic acid (TCA) cycle and therefore impaired respiration. To test this hypothesis, we tested whether overexpression of NOX might similarly rescue virulence defects in *pdhC*::Tn mutants. To test this, we conjugated a NOX overexpression plasmid into the *pdhC::Tn* background and evaluated virulence using the L2 fibroblast plaquing assay, which had previously shown the greatest residual growth in *pdhC::Tn* and provided the least stringent defects for rescue. In contrast to our hypothesis, the *pdhC::Tn* + NOX strain showed no improvement in plaque formation relative to the isogenic *pdhC::Tn* mutant, and remained significantly attenuated compared to WT *L. monocytogenes* (**Supplemental Figure 1A**). This was further confirmed by analysis of fermentative byproducts produced. We found that the *pdhC*::Tn-NOX strain showed modest rescue of acetate production, but not nearly to the extent of menaquinone deficient strains (**Supplemental Figure 1B**). These results suggest that, unlike menaquinone mutants, the virulence defects in *pdhC* mutants cannot be rescued by redox rebalancing through NOX overexpression alone. In sum, we found that rapid regeneration of NAD+ via NOX is insufficient to rescue *pdhC::*Tn virulence in plaquing assays and that the defect must not be driven purely by an inability to regenerate NAD+ via oxidative metabolism.

### Unbiased metabolomics reveals elevated pyruvate and lactate levels in *pdhC*::Tn, but otherwise globally decreased metabolites

Given that the *pdhC::Tn* mutant could not be rescued by NOX overexpression—despite exhibiting an altered respiro-fermentative metabolic profile—we sought to gain a more global view of metabolic dysregulation in this strain. We hypothesized that PDH-deficient strains would accumulate upstream glycolytic intermediates due to impaired flux through pyruvate, while tricarboxylic acid (TCA) cycle intermediates would be depleted owing to inefficient conversion of pyruvate to acetyl-CoA. To test this hypothesis, we performed untargeted metabolomics on WT *L. monocytogenes* and *pdhC*::Tn mutants grown in defined medium supplemented with 110 mM of glucose. The use of defined medium was essential, as complex media, such as BHI, contains abundant background metabolites that interfere with LC/MS metabolite detection and analysis. Additionally, excess glucose supplementation (110 mM versus 55 mM) was required because our lab had observed that PDH mutants show impaired growth in defined medium containing standard glucose concentrations (55mM) (Data not shown). We focused our analysis on glycolytic and TCA cycle intermediates, with supplemental analysis of intracellular levels of the fermentative byproduct lactate. Consistent with our hypothesis, relative to WT *L. monocytogenes*, *pdhC::Tn* mutants exhibited elevated concentrations of pyruvate and lactate, and a marked depletion of TCA cycle metabolites (**Figure 3**). In contrast to our hypothesis however, we observed significantly reduced levels of upper glycolytic intermediates in the *pdhC::Tn* mutants relative to WT (**Figure 3**). Taken together this data suggested three different facets of *pdhC*::Tn mutant metabolism: 1. *pdhC*::Tn was fermenting what sugars it was able to acquire more toward lactate compared to WT. 2. *pdhC*::Tn mutants are reduced in their capacity to funnel metabolites into the TCA cycle compared to WT. 3. *pdhC*::Tn mutants are defective for the acquisition of carbon as represented by lower levels of upper glycolytic metabolites compared to WT.

**Figure 3.**
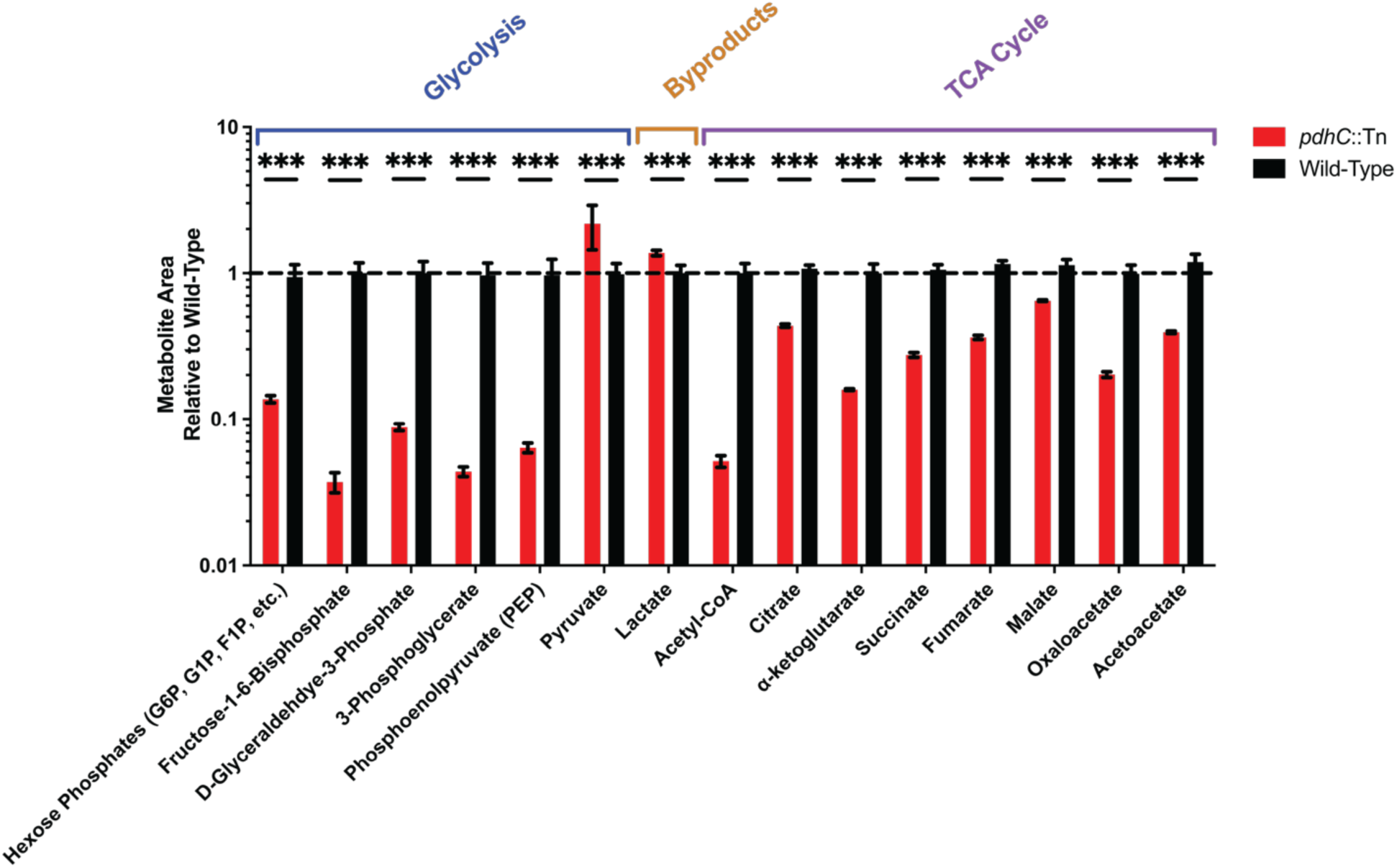
Metabolomic profiling of *pdhC*::Tn and WT *L. monocytogenes* glycolytic, fermentative byproduct, and TCA cycles metabolites. Indicated *L. monocytogenes* strains were grown to mid-log phase (OD_600_ ≈ 0.4) in Listeria Synthetic Medium (LSM) supplemented with 110 mM glucose. Intracellular metabolites were extracted and analyzed via HPLC-MS. Tricarboxylic acid (TCA) cycle intermediates, upper glycolytic metabolites, and lactate were identified based on accurate mass-to-charge (m/z) ratios and retention times using reference values from the KEGG Compound Database, as implemented in MAVEN software. Peak areas were quantified and normalized to wild-type (WT) levels for each metabolite. Data represent three biological replicates, each with two technical replicates. Statistical comparisons were performed using unpaired two-tailed Student’s *t*-tests for each metabolite within a strain.

### *pdhC*:Tn shows an impaired ability to grow in LSM supplied with PTS-mediated carbon sources and can be rescued for growth on PTS-independent hexose phosphates

The unexpected finding that *pdhC::Tn L. monocytogenes* mutants have reduced levels of upper glycolytic intermediates, despite a block downstream in the conversion of pyruvate to acetyl-CoA, led us to hypothesize that *pdhC::Tn* mutants may be defective in acquiring carbon sources, some of which might be available within the host cytosol. This hypothesis was supported by prior observations from our lab showing that *pdhC::Tn* mutants are unable to grow in defined media supplemented with standard concentrations of glucose (55 mM) (Data not shown). Glucose uptake in *L. monocytogenes* occurs primarily through the phosphotransferase system (PTS), in addition to PTS independent GLUC transporters (30,61). This phenotype, coupled with the buildup of pyruvate in *pdhC::Tn* mutants, which would inhibit the PTS dependent phosphor-relay initiated by the conversion of phospoenolpyruvate to pyruvate by EI (*ptsI*), led to the hypothesis that *pdhC::Tn* mutants would show impaired respiration of PTS-mediated carbon sources relative to WT *L. monocytogenes* (32).

To test the hypothesis that *pdhC::Tn* is impaired in its ability to consume PTS-mediated carbon sources we assessed growth in *Listeria* synthetic media (LSM) with glucose, fructose, mannose, and glucoe-6-phosphate (+glutathione) as the sole carbon source (110 mM). In short, LSM containing defined sole carbon sources was inoculated with WT *L. monocytogenes* or the indicated mutants, and growth was monitored by measuring OD_600_ every 15 minutes for 24 hours. Notably, LSM supplemented with hexose phosphates required the addition of 10 mM reduced glutathione to induce *prfA*—and consequently *uhpT*—expression. As expected, WT *L. monocytogenes* was able to grow readily on all of these carbon sources (**Figure 4A-D**). Of note, we found that *pdhC::Tn* was significantly impaired for growth on PTS-mediated carbon sources of glucose, fructose, and mannose (**Figure 4A-C**). This was characterized by slow growth that did not reach the OD600 of WT until nearly 24 hours after inoculation. Importantly, each of these growth defects could be rescued to WT levels with heterologous overexpression of *pdhC*. Interestingly, *pdhC::Tn* showed WT levels of growth in LSM supplied with hexose phosphates (+glutathione), consistent with a specific defect in the acquisition of PTS dependent carbon sources (**Figure 4D**). Taken together, these data demonstrate the *pdhC::Tn* is able to grow on PTS mediated carbon sources but that growth is significantly impaired relative to WT *L. monocytogenes*. Further, *pdhC::Tn* can be rescued for growth in LSM on PTS-independent carbon sources such as hexose phosphates (**Figure 4A-D**).

**Figure 4.**
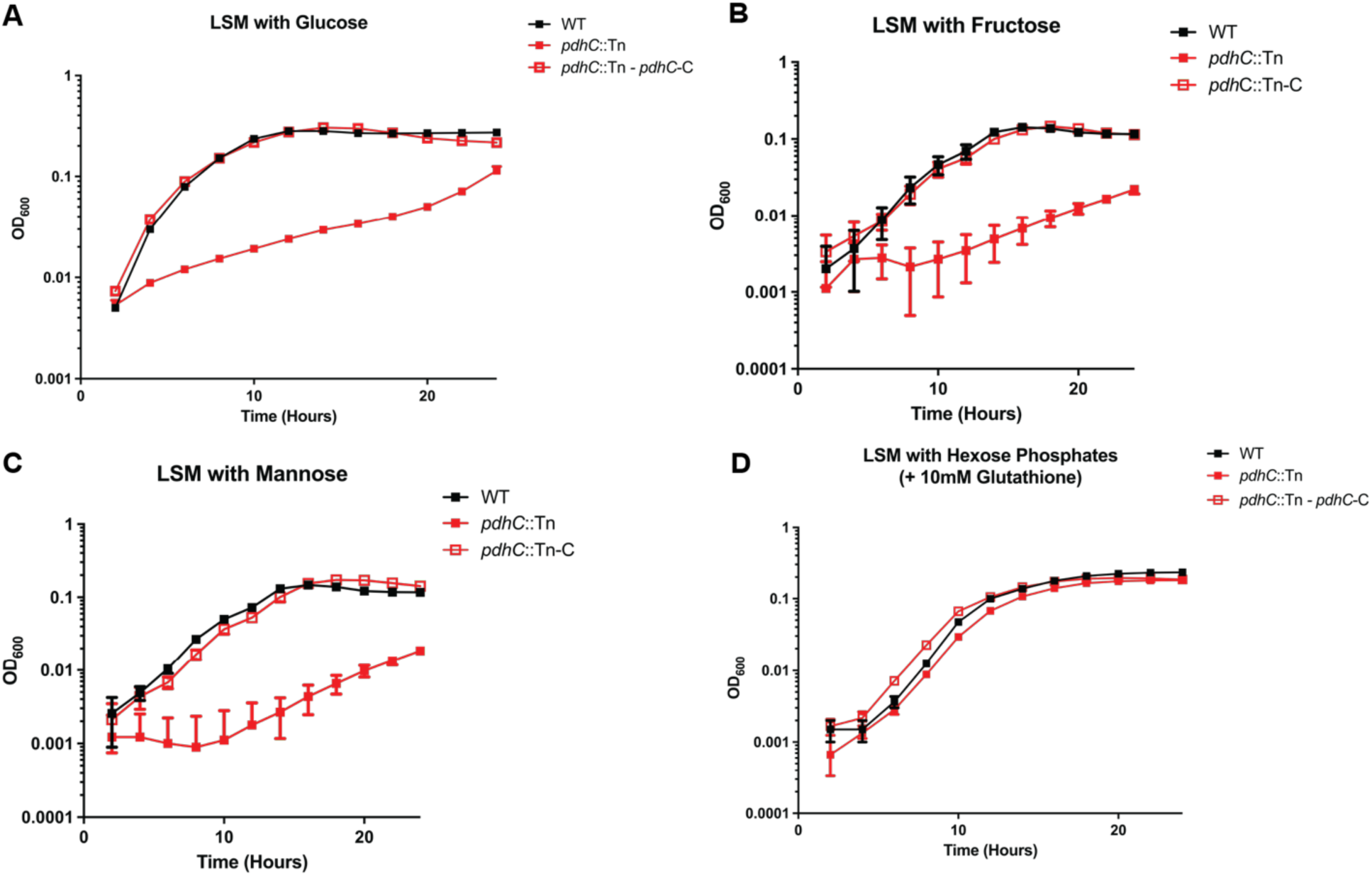
*PdhC*::Tn mutants are defective for growth on PTS-mediated carbon Source of glucose, fructose, and mannose, but retain growth on non-PTS-mediated hexose phosphates. Indicated strains were grown in LSM at 37°C, shaking at 250 r.p.m. with the addition of 110mM glucose (A) or molar equivalent amounts of fructose (B), mannose (C), or hexose phosphates (+10mM glutathione) (D). OD600 was monitored every 15 minutes for 24 hours is a plate reader. Data represents average of three technical replicates from one representative of three biological replicates.

### *PdhC*::Tn suppressor screen reveals strains with restored growth on PTS-mediated carbon sources

*L. monocytogenes* uses host-derived PTS-mediated carbon sources to be able to survive and replicate in the host cytosol (29). As PDH mutants are unable to grow in defined media supplemented with phosphotransferase system (PTS)-mediated carbon sources we hypothesized that a key contributor to the attenuated virulence of PDH mutants is their inability to utilize PTS-dependent carbon substrates (29). To test this hypothesis, we conducted a *pdhC::Tn* mutant suppressor screen on LSM plates supplemented with 55 mM fructose as the carbon source to identify suppressor mutations that would allow growth of PDH deficient mutants on PTS substrates. Fructose was chosen because it is acquired exclusively via the PTS and, as a hexose sugar, provides a carbon input comparable to glucose or glucose-6-phosphate. Additionally, WT *L. monocytogenes* grows robustly in LSM + 55 mM fructose, whereas *pdhC::Tn* mutants require at least 110 mM fructose to support even limited growth (**Figure 4B**). To efficiently induce suppressor mutations, exponentially growing *pdhC::Tn* cultures were mutagenized with ethyl methanesulfonate (EMS) for five minutes, washed to remove residual mutagen, and stored in 40% glycerol at –80 °C. To avoid false-positive suppressors arising from growth on carryover metabolites in frozen stocks, the mutagenized library was thawed, washed with phosphate-buffered saline (PBS), centrifuged, and resuspended in PBS prior to screening. Approximately 10⁷ EMS-mutagenized *pdhC::Tn* mutants were plated per dish across ten LSM + 55 mM fructose plates and incubated at 37 °C for two days. Resulting suppressor colonies were restreaked on selective media to confirm retention of the transposon. Five suppressor colonies that retained the transposon and exhibited restored growth on LSM + 55 mM fructose plates were selected for analysis by whole-genome sequencing. Strikingly, all five suppressor mutants contained mutations in a single gene, *LMRG_01223* - annotated as *rex*, which encodes a redox-sensing transcriptional repressor (**Table 1**). Of the five identified mutations in *rex*, three were missense mutations, one was a premature stop codon, and one was a large C-terminal deletion extending beyond the native stop codon (**Table 1**). Taken together, these results suggest that inactivation of *rex* is a highly reproducible mechanism of restoring growth of *pdhC::Tn* mutants on PTS-dependent fructose.

**Table 1.**
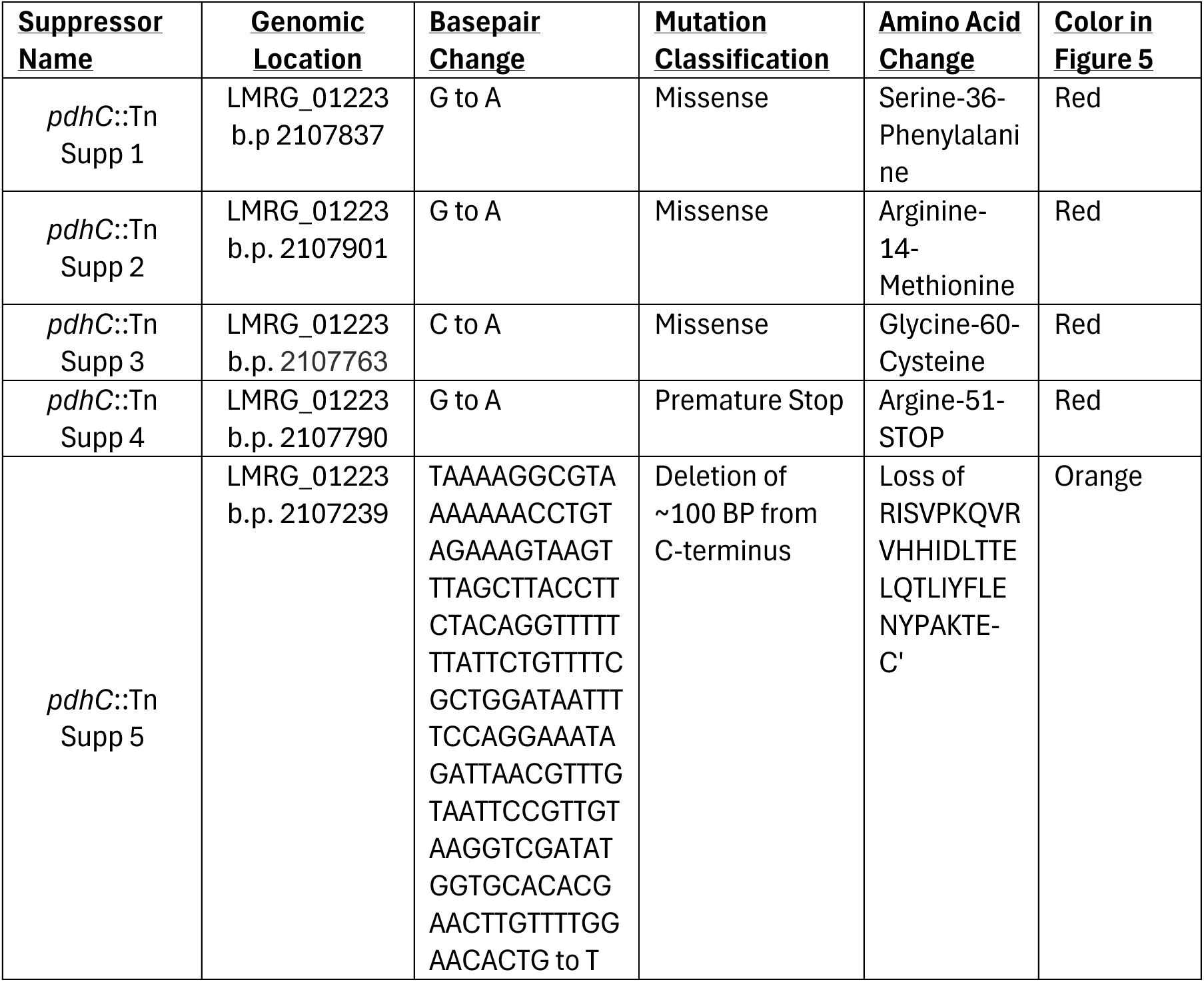
Suppressors mutations in *rex* (LMRG_01223) of *pdhC*::Tn *L. monocytogenes* growth on LSM with fructose plates.

### *pdhC*::Tn suppressor mutations for growth on PTS-mediated carbon sources are primarily in the DNA binding domain of Rex

To better understand how the identified mutations may impact Rex function, we sought to map their locations onto a model of the *L. monocytogenes* Rex structure. Previous studies in other organisms have shown that Rex forms homodimers capable of binding NAD⁺ or NADH, adopting open or closed conformations, respectively (54). In the open conformation, Rex bound to NAD⁺ associates with Rex-specific operator sequences in the bacterial genome to repress genes involved in fermentative metabolism (38,54). This NAD⁺-dependent binding is thought to signal active oxidative metabolism, indicating sufficient NAD⁺ regeneration relative to NADH (54). Consequently, when possible, the bacterium prioritizes respiration over fermentation to maximize ATP production and biosynthetic efficiency (25). To model this interaction, we used AlphaFold3 to predict the structure of the *L. monocytogenes* Rex homodimer in complex with NAD⁺ and target DNA (62). The resulting model was visualized using Jmol (Jmol: an open-source Java viewer for chemical structures in 3D. http://www.jmol.org/), where residue coloring and mutation annotations were added for clarity. Structural modeling revealed that all three missense mutations reside within the predicted DNA-binding domain (alpha helices 1-4) of Rex in its NAD⁺-bound, open conformation (**Figure 5**). This suggests that the mutations likely impair DNA binding and thus prevent Rex from properly regulating gene expression. Notably, none of the mutations are located within the NAD⁺/NADH binding pocket, further supporting the interpretation that impaired DNA binding, rather than cofactor recognition, underlies the observed regulatory defects (**Figure 5**).

**Figure 5.**
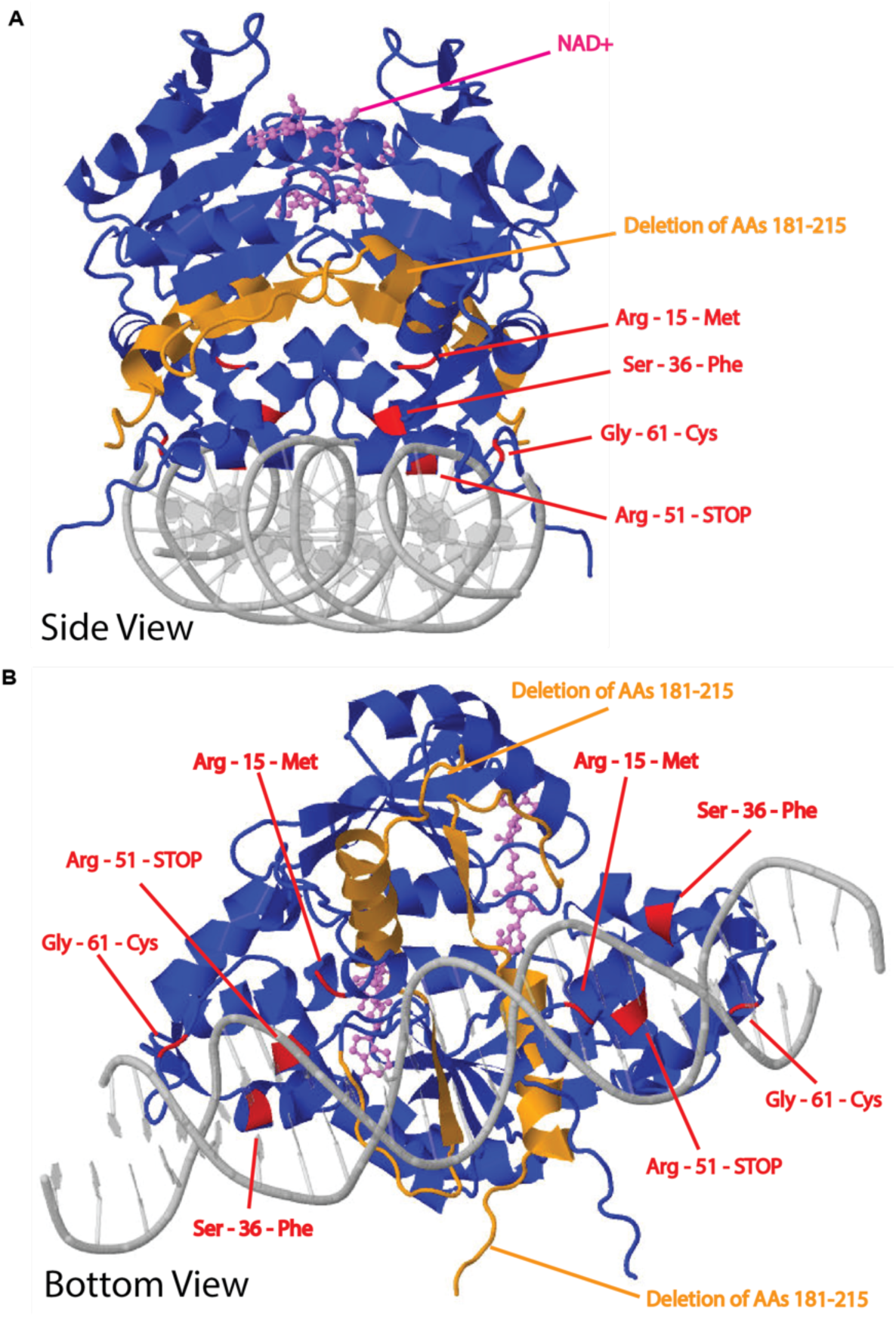
Suppressors of *pdhC*::Tn growth on PTS mediated carbon sources mapped onto *L. monocytogenes*’ Rex (LMRG_01223) homodimer bound to NAD+ and DNA. *L. monocytogenes (10403s)* protein sequence for Rex (LMRG_01223) was obtained from NCBI (GCA_000168695.2_ASM16869v2) and was input into AlphaFold as a homodimer with the ligands of NAD+ (pink) and Rex-specific DNA-binding sequences (grey DNA helix). Predicted output was further processed in Jmol to represent amino acids modified (red) or lost (orange) due to suppressor mutations. Complete molecular structure is pictured from two angles: side-side (A) and bottom-up (B).

### *pdhC*::Tn suppressor mutants show restored growth in LSM with fructose

Suppressor mutations enabling *pdhC::Tn* growth on LSM supplemented with fructose were initially identified on solid media but not in liquid cultures. To confirm that these mutations would similarly support growth on PTS-mediated carbon sources in liquid media, we assessed growth in liquid LSM containing fructose. As expected, WT *L. monocytogenes* exhibited robust growth in LSM+fructose, while the *pdhC::Tn* mutant displayed severely impaired growth, requiring approximately 24 hours to begin approaching the terminal OD_600_ achieved by WT (**Figure 6**). Notably, all five suppressor mutants demonstrated rescued growth in LSM with fructose, reaching OD_600_ values comparable to WT with similar kinetics (**Figure 6**). These results validate that the identified suppressor mutations permit *pdhC::Tn* to grow efficiently on fructose in both solid and liquid media, supporting the conclusion that loss of *rex*-mediated repression facilitates PTS-dependent carbon source utilization.

**Figure 6.**
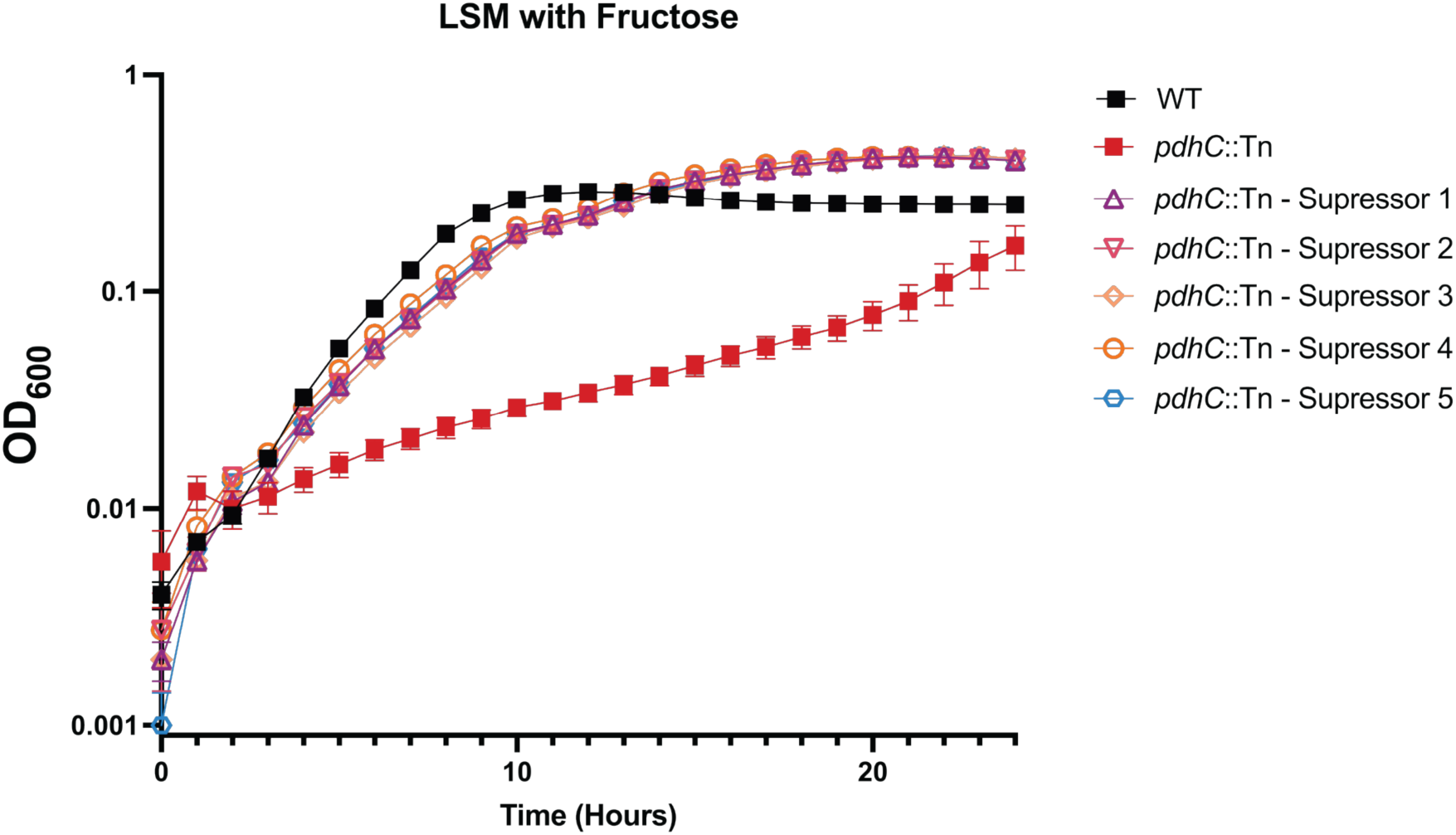
*PdhC*::Tn suppressor mutants restore growth in define media with fructose as the sole carbon source. Indicated strains were grown in LSM at 37°C, shaking at 250 r.p.m. with the addition of 110mM Fructose. OD600 was monitored every 15 minutes for 24 hours is a plate reader. Data represents average of three technical replicates from two biological replicate.

### *pdhC*::Tn mutants cannot grow on PTS-mediated carbon sources in oxygenated defined media, but can when grown anaerobically or on PTS-independent carbon sources

Previous work by Halsey et al. (2021) demonstrated that *L. monocytogenes* Rex functions as a transcriptional repressor of fermentative metabolism when respiration is available, a state sensed through elevated NAD⁺ levels (38). Building upon this finding—and considering that all five identified *rex* mutations in our suppressor screen either disrupt the predicted DNA-binding domain or result in presumed loss-of-function alleles (e.g., premature stop codons or large C-terminal deletions)—we hypothesized that loss of Rex activity in the *pdhC::Tn* background relieves fermentative repression (**Table 1 and Figure 6**). If true, *pdhC::Tn* mutant growth should be rescued on fructose under anaerobic conditions and should phenocopy the *pdhC::Tn-rex* suppressor mutants grown under aerobic conditions. To test this hypothesis, we compared the growth of WT, *pdhC::Tn*, *pdhC::Tn* complemented with *pdhC* (*pdhC::Tn* + *pdhC-C*), and *pdhC::Tn* Suppressor #4 (Rex – Arg51-STOP) under both aerobic and anaerobic conditions. Cultures were inoculated into LSM containing 110 mM fructose and incubated at 30 °C for 48 hours, either aerobically (stationary incubation) or anaerobically in a GasPak chamber. After incubation, cultures were mixed and transferred to a 96-well plate for optical density measurement at 600 nm (OD_600_). Under aerobic conditions, WT, *pdhC::Tn* + *pdhC-C*, and *pdhC::Tn* Suppressor #4 (Rex – Arg51-STOP) all achieved comparable levels of growth (**Figure 7A**). In contrast, in aerobic conditions *pdhC::Tn* exhibited markedly impaired growth, supporting the interpretation that functional Rex represses growth on fructose when aerobic respiration is possible (**Figure 7A**). Under anaerobic conditions, however, all strains—including *pdhC::Tn*—grew to similar levels, consistent with the loss of Rex-mediated repression in the absence of respiration (**Figure 7B**). Taken together, these findings support the conclusion that *pdhC::Tn* is capable of growing on PTS-mediated fructose in defined media either by genetic disruption of *rex* or by shifting to anaerobic culture conditions that naturally relieve fermentative repression.

**Figure 7.**
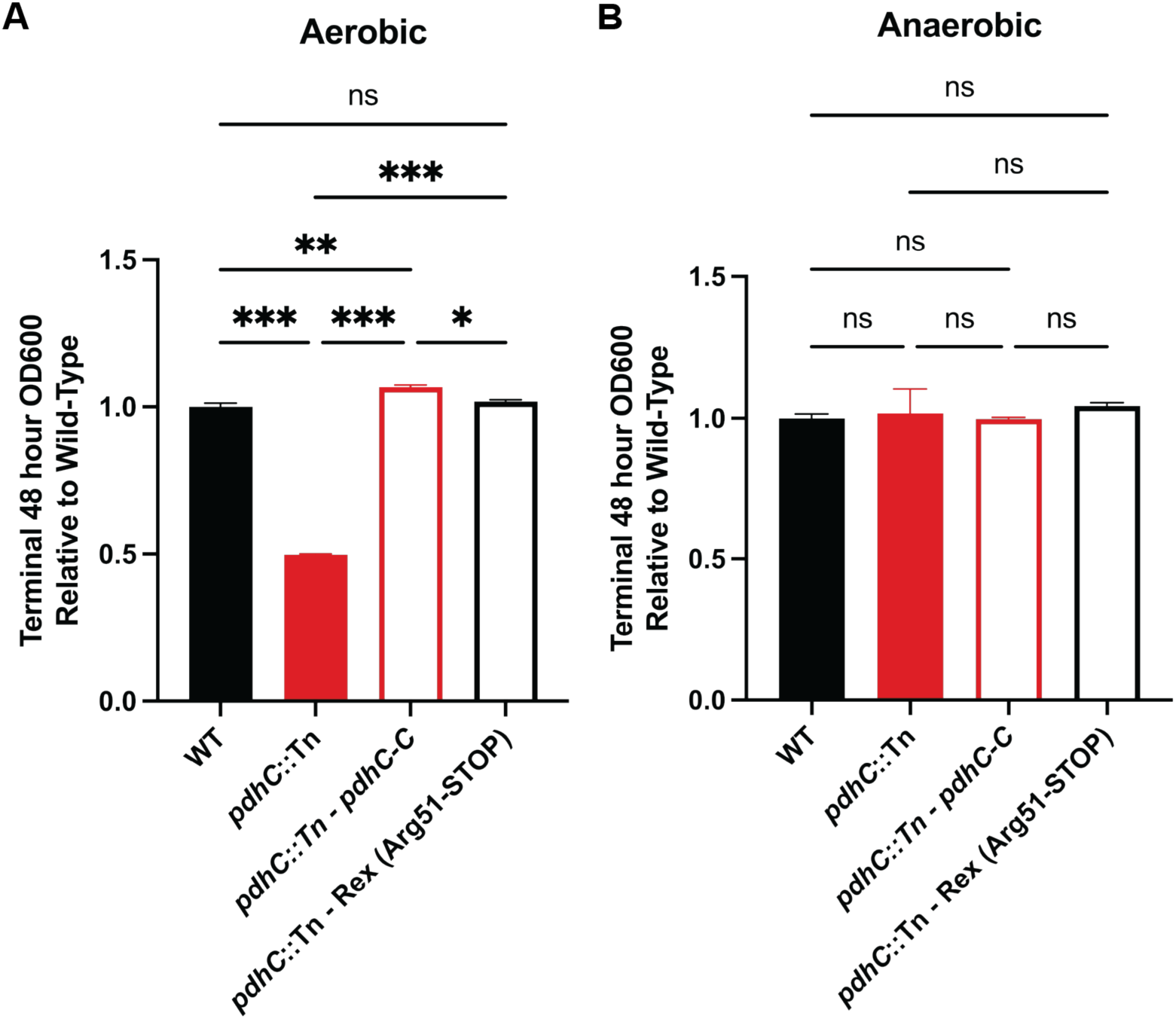
Loss of rex permits *pdhC*::Tn growth on PTS-mediated carbon sources aerobically similar to that of anaerobic growth. Indicated strains were grown in LSM at 30°C, stationary for 48 hours with the addition of 110mM Fructose. OD600 was taken at 48 hours and normalized to WT. Data represents average of three biological replicates and statistical analysis was performed comparing all strains using one-way ANOVA with Tukey correction.

### *pdhC*::Tn supressor #4 (Rex – Arg51-STOP) rescues intramacrophage growth, but not *in vivo* virulence

Our lab has previously demonstrated that *Listeria monocytogenes* requires a functional phosphotransferase system (PTS) for growth within the macrophage cytosol and for full virulence *in vivo* (29). We hypothesized that *rex* suppressor mutants of *pdhC::Tn* that rescue PTS-mediated growth would partially restore virulence. To test this hypothesis, we assessed the ability of *pdhC::Tn* Suppressor #4 (Rex – Arg51-STOP) to replicate within the host cytosol of BMDMs. As expected, WT *L. monocytogenes* grew robustly in the macrophage cytosol, expanding by approximately 1.5 logs over an 8-hour infection (**Figure 8A**). In contrast, *pdhC::Tn* mutants failed to grow and were progressively cleared, consistent with prior observations (**Figure 8A and Figure 1B**). Remarkably, *pdhC::Tn* Suppressor #4 (Rex – Arg51-STOP) mutants exhibited intracellular replication comparable to WT, indicating that loss of Rex-mediated fermentative repression not only restored growth on PTS carbon sources *in vitro* but also enabled survival and replication in the macrophage cytosol (**Figure 8A**). These data suggest that inability to acquire PTS-dependent carbon sources restricts survival and replication of PDH deficient mutants in the macrophage cytosol.

**Figure 8.**
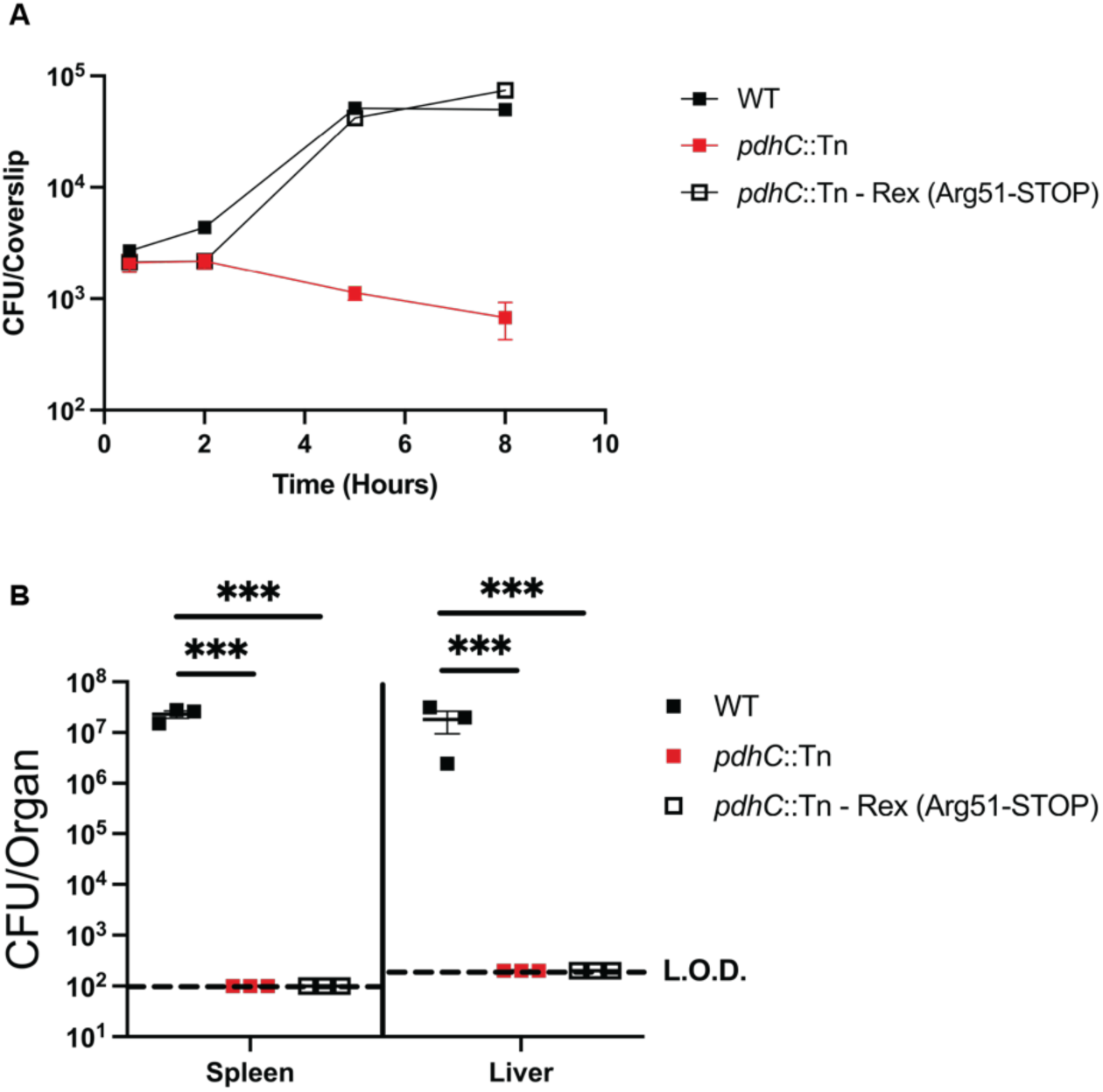
*PdhC*::Tn rex suppressor mutations rescue intramacrophage growth but fail to rescue in vivo. (A) Intracellular growth of indicated strains was determined in BMDMs following infection at an MOI of 0.2. Growth curves are representative of 3 independent experiments. Error bars represent the standard error of the means of technical triplicates within the representative experiment. (D) Bacterial burdens from the spleen and liver were enumerated at 48 hours post-intravenous infection with 1×105 bacteria. Data are representative of results from one experiment. Horizontal dashed lines represent the limits of detection, and the bars associated with the individual strains represents the mean and SEM of the group.

To evaluate whether this rescue extended to *in vivo* infection, we performed an acute murine virulence assay. C57BL/6 mice were intravenously infected with 10⁵ CFU of each strain suspended in 200 μL of PBS. After 48 hours, mice were euthanized and bacterial burdens in the spleen and liver were enumerated. WT *L. monocytogenes* successfully colonized both organs with bacterial burdens of ∼10^7^ CFU/organ, while *pdhC::Tn* was completely attenuated, with burdens falling below the assay’s limit of detection (**Figure 8B**). In contrast to the rescue of intracellular growth observed in BMDMs, *pdhC::Tn* Suppressor #4 (Rex – Arg51-STOP) did not rescue virulence *in vivo* and exhibited organ burdens indistinguishable from the parental *pdhC::Tn* mutant (**Figure 8B**). These results suggest that, although Rex inactivation permits cytosolic replication in cultured macrophages, additional host-specific pressures *in vivo* prevent the suppressor strain from establishing systemic infection. This implies that the murine host imposes metabolic or immunological constraints more stringent than those encountered in *ex vivo* macrophage models—constraints that remain restrictive for both *pdhC::Tn* and its *rex* suppressor derivative.

## DISCUSSION

Mechanisms of carbon acquisition, catabolism, and anabolism are critical virulence determinants that support the pathogenesis of intracellular bacterial pathogens (29,36,41,42). However, these mechanisms remain incompletely defined. In particular, how bacterial pathogens acquire nutrients and efficiently metabolize these nutrients while evading host defenses within the nutrient-limited and hostile environment of the cytosol is not fully understood (17). Elucidating how pathogens regulate metabolism during infection is essential to identifying key metabolic determinants of virulence and potential targets for antimicrobial therapies (63). *Listeria monocytogenes* is both an important human pathogen and a well-characterized model organism (3,64). Despite extensive study, our understanding of the metabolic factors that contribute to its virulence remain limited. In this work, we expand upon findings from Chen et al. (2018), which demonstrated that a *pdhC::Tn* mutant of *L. monocytogenes* exhibits a severe defect in cytosolic growth within macrophages and near-complete attenuation of virulence (20). To further investigate the role of the pyruvate dehydrogenase (PDH) complex during infection, we characterized additional transposon mutants in two other subunits of this complex: *pdhA::Tn* and *pdhD::Tn*. *pdhA::Tn* and *pdhD::Tn* mutants phenocopy *pdhC::Tn*— exhibiting WT growth in rich media but defective intracellular replication and virulence in both L2 fibroblast plaquing assays and acute murine infection models. We then focused on dissecting the virulence defect of *pdhC::Tn* and found that it has altered respiro-fermentative metabolism, producing elevated levels of lactate and reduced levels of acetate compared to WT *L. monocytogenes*. This secreted metabolic profile resembles that of previously described *L. monocytogenes* mutants impaired in respiration (21–23). However, unlike those mutants, the virulence defect of *pdhC::Tn* was not rescued by heterologous expression of NADH oxidase (NOX). Unbiased metabolomic analysis further revealed that *pdhC::Tn* accumulates pyruvate and lactate while being depleted in upper glycolytic and tricarboxylic acid (TCA) cycle intermediates. Using targeted growth experiments of *pdhC*::Tn in defined media with PTS-dependent and -independent carbon sources, we discovered that *pdhC::Tn* is significantly impaired in its ability to utilize phosphotransferase system (PTS)-mediated carbon sources, but can grow readily on the PTS-independent carbon source of hexose phosphates. Based on this observation, we performed a suppressor screen to select for mutants capable of growing on minimal medium containing fructose as the sole PTS-dependent carbon source. Whole-genome sequencing and SNP analysis of five independent suppressor strains revealed mutations in a single gene, *rex* (*LMRG_01223*), a redox-sensing transcriptional repressor. These mutations included missense mutations in the DNA-binding domain, premature stop codons, and truncations likely resulting in loss of function. Functional assays demonstrated that *rex* acts as a fermentative repressor in *pdhC::Tn* mutants grown on PTS-dependent carbon sources and that loss of *rex* in a *pdhC::Tn* background restored intracellular growth in macrophages ex vivo but failed to rescue virulence in murine infection models. These findings further support that *L. monocytogenes* requires the ability to acquire and use PTS-mediated carbon sources to grow in the host cytosol, but also unveil that other pleiotropic defects of PDH complex mutants must be critical for full virulence *in vivo.* Furthermore, they indicate that Rex likely promotes respiratory metabolism in the host cytosol through fermentative repression, which is essential for full virulence (23).

Although central glycolytic and TCA cycle enzymes have been extensively studied for their roles in carbon metabolism, their contributions to bacterial pathogenesis remain underexplored across many species (25,65). While the necessity of the PDH complex during *L. monocytogenes* infection is loosely established, the specific physiological mechanisms leading to the avirulent phenotype of PDH mutants—despite their robust growth in rich media—has not been elucidated (18). Interestingly, all PDH subunit mutants (*pdhA::Tn*, *pdhC::Tn*, and *pdhD::Tn*) retain the capacity to form plaques in L2 fibroblast monolayers, albeit significantly smaller than WT, suggesting that while the PDH complex is important across multiple host environments, it is not universally essential. This phenotypic discrepancy points to differences in the pressures imposed by distinct host cell types, experimental timelines, or experimental conditions. One hypothesis is that plaque formation may occur independently of robust intracellular replication. Preliminary evidence suggests that *L. monocytogenes* can spread cell-to-cell when host cells reach their carrying capacity, even under metabolically limiting conditions (66). Thus, PDH mutants might form plaques despite failing to survive within macrophages or establish infection *in vivo* due to the ability to bypass metabolic depletion. Differences in the ability of PDH complex mutants to grow in L2 fibroblasts, but not macrophages, may also reflect variation in host cell detection and response to altered bacterial metabolism, possibly driven by differential nutrient availability or immune signaling thresholds. It has been shown that intracellular levels of lactate and acetate can act as signal of infection and host cell response (67–70). Although not addressed in this study, we hypothesize that individual PDH subunit mutants altered fermentative byproducts may be impacting interaction with L2 fibroblast and macrophages, divergently. One way to further identify how production of these organic acids impact host response would be to limit the ability of PDH complex mutants to produce them through deletion of acetate kinase and/or lactate dehydrogenases and assess virulence capabilities and host cell responses.

One key unanswered question in the field is the relative contribution of fermentative versus respiratory metabolism during infection. *L. monocytogenes* appears to balance these states, potentially as an evolutionary adaptation to evade host detection of metabolic byproducts. Rex, previously characterized by Halsey et al. (2021), is largely dispensable for virulence in WT strains (38). However, its role in repressing fermentation during infection may be context-dependent. For instance, *aro* mutants and menaquinone-deficient mutants lacking respiratory capacity are highly attenuated, but both have only been assessed in the presence of Rex (21–23,60). Our findings suggest that Rex enforces respiratory metabolism in macrophages, and loss of Rex may permit survival via fermentation and restoration of PTS-mediate carbon source use. To dissect the respective roles of fermentation and respiration, strains lacking essential respiratory components should be tested in both *rex*-positive and *rex*-negative contexts.

Conversely, strains deficient in fermentation could be constructed with Rex overexpression to assess reliance on respiratory metabolism. An example of this would be to delete *ackA* and *ldh*, enzymes essential for *L. monocytogenes’* production of fermentative byproducts with Rex hyperexpression. This mutant would in theory be completely dependent on respiration for growth and could isolate one side of the respiro-fermentative metabolism. These experiments will clarify how host cells detect and respond to distinct bacterial metabolic states and what is whether respiration or fermentation is the predominant need of cytosolic pathogens.

Unexpectedly in *pdhC::Tn*, acetate is still predominantly produced despite the absence of a functional PDH complex. The suggests that *pdhC*::Tn can still funnel substantial amounts of carbon into acetyl-CoA and the TCA cycle, supporting a minimal respiratory capacity. We hypothesize that some of this metabolic flux is mediated by alternative pathways of pyruvate metabolism include pyruvate oxidase and pyruvate carboxylase (25). It would seem unlikely this flux is mediated via enzymes such as pyruvate formate lyase, which are intoxicated in the presence of oxygen due to a glycyl free-radical (71). Nevertheless, the conversion of pyruvate to acetyl-CoA, and therefore acetate, seems to be critical for full virulence of *L.* monocytogenes. This is supported by the fact that neither loss of *rex* or addition of NOX was sufficient to rescue *pdhC*::Tn. Understanding what pathways are active for the conversion flux into the TCA cycle may be critical to understanding *L. monocytogenes* virulence in the cytosol.

Another question raised by this work is the role of *rex* in *L. monocytogenes’* metabolic regulation and its role during *L. monocytogenes* pathogenesis. Preliminary work to identify how *rex* impacts virulence has been previously characterized by Halsey *et al*. 2021 (38). In which, they showed that *rex* mutants replicate readily in the macrophage cytosol, form larger plaques than WT *L. monocytogenes*, and are only modestly attenuated for *in vivo* virulence of the liver and spleen via an oral infection model (38). Importantly, to date nobody has assessed a *rex* mutant during intravenous infections, which we hypothesize may show less attenuation via this method of delivery due lack of metabolic transition from the gut to intracellular environment. However, it is also possible that this strain could be more attenuated due to *rex* mutants lacking the ability to repress their fermentation and therefore having a metabolism not well-adjusted for the *in vivo* cytosolic environment. In either case, it is difficult to assess what the metabolism of a *rex* mutant is during cytosolic growth. This is obfuscated by the fact that *rex* mutants don’t lack any metabolic capabilities, rather they have access to metabolic pathways likely repressed during infection. Therefore, it is important to ask why certain metabolic pathways are in fact repressed by *rex* during infection and does relying on these pathways for energy generation impact virulence. Initial work done by the Reniere lab has evaluated genes known to be regulated by *rex* and they have shown a wide variety of genes repressed including many PTS, core fermentative enzymes like lactate dehydrogenase and pyruvate format lyase, and some virulence genes including internalins A and B (38). One important question raised by these findings is how over expression of some of these enzymes may independently impact PDH mutants’ virulence with *rex* intact, such as lactate dehydrogenase. While restoration of some PTS expression likely explains the phenotypes identified in this work, it is possible that use of PTS could be independently impacting virulence due to the intertwined nature of metabolism and virulence gene regulation. While it is clear that *rex* participates in the repression of internalins, it is possible that *rex*-mediated repression of PTS may be further impacting *prfA* and other virulence gene expression. For that reason, it would be important to assess whether PrfA* addition can further rescue virulence phenotypes of *rex* and PDH mutants. Together, this work could further define how *pdhC*::Tn is rescued by loss of *rex* and further how metabolic shifts *in vivo* may be connected to virulence gene expression.

Pinpointing the exact cause of virulence defects in PDH mutants is challenging due to the centrality of this enzymatic complex in metabolism. While we show that carbon acquisition via PTS is impaired in PDH-deficient strains, additional physiological perturbations likely contribute to its attenuation. Notably, cell-type-specific phenotypes suggest that individual host environments present distinct metabolic challenges or immune barriers. The restoration of intramacrophage growth in *rex*-deficient *pdhC::Tn* mutants, coupled with persistent *in vivo* attenuation, implies that host tissues may be more metabolically stringent or better equipped to detect altered bacterial metabolism. It is possible that PDH complex mutants with loss of function Rex may be producing excess lactate and this is being detected by host cells *in vivo,* but not *ex vivo*. Potential reasons for lack of *ex vivo* detection could be the supraphysiologic conditions as well as pH buffering of the media. A deeper understanding of how host cells detect bacterial fermentation versus respiration will yield insight into both immune surveillance mechanisms and pathogen evasion strategies. Similarly, understanding how *pdhC::Tn* retains the ability to bypass PDH complex mediated conversion of pyruvate into acetyl-CoA may unveil pathways essential for virulence.

## MATERIALS AND METHODS

### Ethics Statement

All animal-based experiments were performed using the protocol (M005916-R01-A01) approved by the Animal Use and Care Committee of the University of Wisconsin—Madison and consistent with the standards of the National Institutes of Health.

### Bacterial Strains and Culture

All *Listeria monocytogenes* strains used for experiments in this study were in a 10403s background. All *L. monocytogenes* strains were grown overnight in BHI and at 30°C stationary for all experiments, except as described. *Escherichia coli* strains were grown in Luria broth (LB) at 37°C shaking. Antibiotics used on *E. coli* were at a concentration of 100 µg/ml carbenicillin or 30 µg/ml kanamycin when appropriate. Antibiotics used on *L. monocytogenes* were at a concentration of 200 μg/mL streptomycin and/or 10 μg/mL chloramphenicol and/or 2 μg/mL erythromycin, when appropriate. Plasmids were transformed into chemically competent *E. coli* and further conjugated in *L. monocytogenes* using S17 *E.coli*.

### Construction of Strains

The pPL2 integrative vector pIMK2 was used for constitutive expression of *L. monocytogenes* genes (72). pIMK2 complement constructs were cloned in XL1-Blue *E. coli* with 30μg/mL Kanamycin and grown for plasmid harvest using Promega MiniPrep Kit. Harvested plasmid sequences were confirmed using was performed by Plasmidsaurus using Oxford Nanopore Technology with custom analysis and annotation. Plasmid were then shuttled into *L. monocytogenes* through conjugation with S17 (pIMK2) *E. coli*. All mutants were confirmed via PCR, plasmid sequencing, and whole-genome sequencing using Oxford Nanopore technology from Plasmidsaurus with custom analysis and annotation.

### *In vitro* Growth Assays

Bacteria were grown overnight in BHI at 30°C stationary. Overnight cultures were used to generate inoculums with ∼3.7×10^8^ CFU in PBS. 100 µLs per well of a flat bottom clear 96-well plate of *Listeria* synthetic media (LSM) with carbons source (supplied with amounts noted in text and figures) was inoculated with 2 µL of inoculums. Plates were parafilmed on the edge to prevent evaporation and evaluated for OD_600_ in plate reader at 37°C shaking (250 r.p.m.) and reads every 15 minutes for times displayed.

### Terminal Optical Density Aerobic and Anaerobic Growth Assays

Bacteria were grown overnight in BHI at 30°C stationary. Overnight cultures were used to generate inoculums with ∼3.7×10^8^ CFU in PBS. 14 mL tubes were setup with 3 mL of *Listeria* synthetic media (LSM) with carbon sources (supplied with amounts noted in text and figures) and inoculated with 20 µL of inoculums. Tubes were loosely capped and placed, slanted, in 30°C incubator either exposed to air (aerobic) or placed in GasPak (anaerobic) chambers with 2 GasPaks for oxygen depletion (Fischer: 11-816-2). Samples were left for 48 hours and then 100 µL was harvested from each and plated into 96-well plate for OD600 to be taken in a plate reader. Optical density values were normalized to WT and averaged for display and statistical analysis.

### Intra-macrophage Growth Curves

Bone marrow-derived macrophages were isolated from CL57/BL6 mice and cultured as previously described in Roswell Park Memorial Institute Medium (RPMI) based media (Invitrogen: 11875093) (73). BMDMs were plated into 60 mm dishes contain 13 degassed coverslips. BMDMs cells were infected with *L. monocytogenes* strains at a multiplicity of infection [MOI] of 0.2. Inoculums of *L. monocytogenes* were grown in 3mL of BHI at 30°C stationary until all strains had reach stationary phase. Colony forming units to OD_600_ ratios were determined for each strain and adjusted to ensure infection results in a comparable MOI across strains. After 30 minutes BMDM media was exchanged for media containing 50 µg/ml Gentamycin. Coverslips were harvested, cells lysed in pure water, bacteria rescued isotonically, and plated to quantify CFU at displayed time points. All strains were assayed in biological triplicate and data displayed is one representative biologic replicate.

### Plaque Assays

Plaque assays were conducted using a L2 fibroblast cell line grown in Dulbecco’s Minimal Essential Media (DMEM) based media (Thermo Fischer: 11965092) as previously described with minor modifications for visualization and quantification of plaques (21). L2 fibroblasts were seeded at 1.2 × 10^6^ per well of a 6-well plate, then infected at an MOI of 0.5 to obtain approximately 10-30 PFU per dish. Inoculums of *L. monocytogenes* were grown in 3mL of BHI at 30°C stationary until all strains had reached stationary phase. Colony forming units to OD_600_ ratios were determined for each strain and adjusted to ensure infection results in a comparable MOI across strains. At 4 days postinfection, cells were stained with 0.3% crystal violet for 10 min and washed twice with deionized water. Stained wells were scanned, uploaded, and areas of plaque formation were measured on ImageJ analysis software. All strains were assayed in biological triplicate and the average plaque areas of each strain (one-well per strain were normalized to wild-type plaque size within each replicate.

### Murine Infection and Organ Burdens

Infections were performed as previously described (21). Briefly, 6 to 12-week-old female and male C57BL/6 mice were infected IV with 1×10^5^ CFU logarithmically growing *L. monocytogenes* (optical density at 600 nm [OD600] = 0.5) in 200 µL of PBS. Colony forming units to OD600 ratios were determined for each strain and adjusted to ensure infection results in a comparable MOI across strains. 48 hours post-infection, mice were euthanized, and livers and spleens were harvested, homogenized in water with 0.1% NP-40, and plated for CFU. Two independent replicates of each experiment with 5 mice per group were performed.

### Fermentation Byproduct Measurements

Indicated strains of *L. monocytogenes* were grown in BHI at 37°C, shaking overnight. Cultures were centrifuged to pellet bacteria, and 1 mL of the supernatant was filtered using a 0.2µm-pore-size syringe filter (09-740-113; Fisher Scientific). Supernatants were then treated with 2μL of H_2_SO_4_ to precipitate running buffer incompatible bacterial components. The samples were then centrifuged at >16000 r.c.f. for 10 min. Subsequently, 200μL of each supernatant was transferred to an HPLC vial. HPLC analysis was performed using a ThermoFisher (Waltham, MA) Ultimate 3000 UHPLC system equipped with a UV detector (210 nm). Compounds were separated on a 250 × 4.6 mm Rezex^©^ ROA-Organic acid LC column (Phenomenex Torrance, CA) run with a flow rate of 0.2 mL min^−1^ and at a column temperature of 50 °C. Prior to injection samples were kept at 4 °C. Separation was isocratic with a mobile phase of HPLC grade water acidified with 0.015 N H_2_SO_4_ (415 µL L^−1^). Byproduct standards were 100, 20, 4, and 0.8mM concentrations of lactate or acetate. HPLC peaks were analyzed and quantified using the Thermofisher Chromeleon 7 software package.

### Metabolic Profiling Using HPLC

Overnight cultures of wild-type (WT) and *pdhC::Tn L. monocytogenes* were grown in brain heart infusion (BHI) broth at 30°C. The following day, 1 mL of each culture was washed with phosphate-buffered saline (PBS) and used to inoculate 50 mL of Listeria Synthetic Medium (LSM) supplemented with 110 mM glucose in baffled flasks. Cultures were incubated at 37°C with shaking until mid-log phase (OD_600_ ≈ 0.4) was reached. At this point, 5 mL of each culture was filtered through a 0.2 μm nylon membrane filter. Filters were then transferred to sterile petri dishes containing 1.5 mL of cold extraction solvent (acetonitrile:methanol:water, 2:2:1). The solvent was gently swirled and pipetted across the filter surface to extract intracellular metabolites, after which the filter was flipped and the process repeated to maximize extraction efficiency. The pooled extract was transferred to centrifuge tubes, vortex vigorously for 2 minutes, and centrifuged at maximum speed (≥13,000 × *g*) for 5 minutes to pellet insoluble material. A 200 μL aliquot of the clarified supernatant was collected, dried under a stream of nitrogen gas, and resuspended in 70 μL of HPLC-grade water prior to analysis. All cultures were grown in biological triplicate and processed in technical duplicate.

Metabolite quantification and analysis was performed as previously described. In short, samples were run through an ACQUITY UPLC BEH C18 column in an 18-min gradient with Solvent A consisting of 97% water, 3% methanol, 10 mM tributylamine (TBA), 9.8 mM acetic acid, pH 8.2, and Solvent B being 100% methanol. Gradient was 5% Solvent B for 2.5 min, gradually increased to 95% Solvent B at 18 min, held at 95% Solvent B until 20.5 min, returned to 5% Solvent B over 0.5 min, and held at 5% Solvent B for the remaining 4 min. Ions were generated by heated electrospray ionization (HESI; negative mode) and quantified by a hybridquadrupole high-resolution mass spectrometer (Q Exactive Orbitrap, Thermo Scientific). MS scans consisted of full MS scanning for 70 to 1,000 m/z from time zero to18 min, except that MOPS m/z of 208 to 210 was excluded from 1.5 to 3 min. Metabolite peaks were identified from Kegg Known Compound list and quantified in Metabolomics Analysis and Visualization Engine (MAVEN).

### *In vitro* Suppressor Screen

The Δ*pdhC*::Tn mutant was mutagenized by a 5-minute exposure to ethyl methanesulfonate (EMS) as previously described (74,75). One ml of the library was thawed, washed in 10 mL PBS and resuspended in PBS to a concentration of 7×10^8^ CFU/mL, and plated across 10 *Listeria* synthetic media (LSM) agar plates with 55 mM fructose. Plates were incubated for 48 hours post-inoculation, at which time single colonies were picked and selected for on BHI plates with erythromycin and streptomycin to confirm resistance. Successful growth of colonies were grown overnight at 37°C with shaking, pelleted by centrifugation, resuspended in 100 μl BHI + 40% glycerol, and stored at -80°C. All five isolates from the LSM with fructose plates was subsequently grown overnight in BHI broth, and subjected to whole genome sequencing and SNP analysis as described below.

### Whole Genome Sequencing and SNP Identification

*PdhC*::Tn *L. monocytogenes* suppressor isolates were grown overnight in 3 mL culture of BHI. Genomic DNA was purified using the MasterPure Gram-positive DNA purification kit (Epicentre) per the manufacturer’s instructions, except that 5 U/μl mutanolysin was used instead of lysozyme. DNA was submitted Plasmidsaurus for whole-genome sequencing using Oxford Nanopore technology. Fastq reads were uploaded to Galaxy and mapped onto the *L. monocytogenes* 10403S reference sequence (GCA_000168695.2_ASM16869v2) using Snippy (version 4.6.0). Single nucleotide polymorphisms (SNPs) were assessed for impact on *L. monocytogenes* coding sequences and genes manually utilizing Jbrowse (Version 1.16.11).

### Cell Culture

L2 cells were all kind gifts from Daniel Portnoy (UC Berkeley). Bone marrow-derived macrophages (BMDM) were prepared from 6-to-8-week-old mice as previously described (76).

### Statistical Analysis

Prism 6 (GraphPad Software) was used for statistical analysis of data. Means from two groups of BioLog plates were compared with unpaired two-tailed Student’s T-test. Means from more than two groups for all other assays were analyzed by one-way ANOVA test. Independently, Mann-Whitney Test was used to analyze two group comparison of non-normal data from animal experiments. * *p* < 0.05, ** *p* < 0.01, *** *p* < 0.001 for all statistical tests displayed.

