## Supplemental Figure 1 and Table 1 for "Understanding the Role of Pyruvate Dehydrogenase in *Listeria monocytogenes* Virulence"

### SUPPLEMENTARY INFORMATION

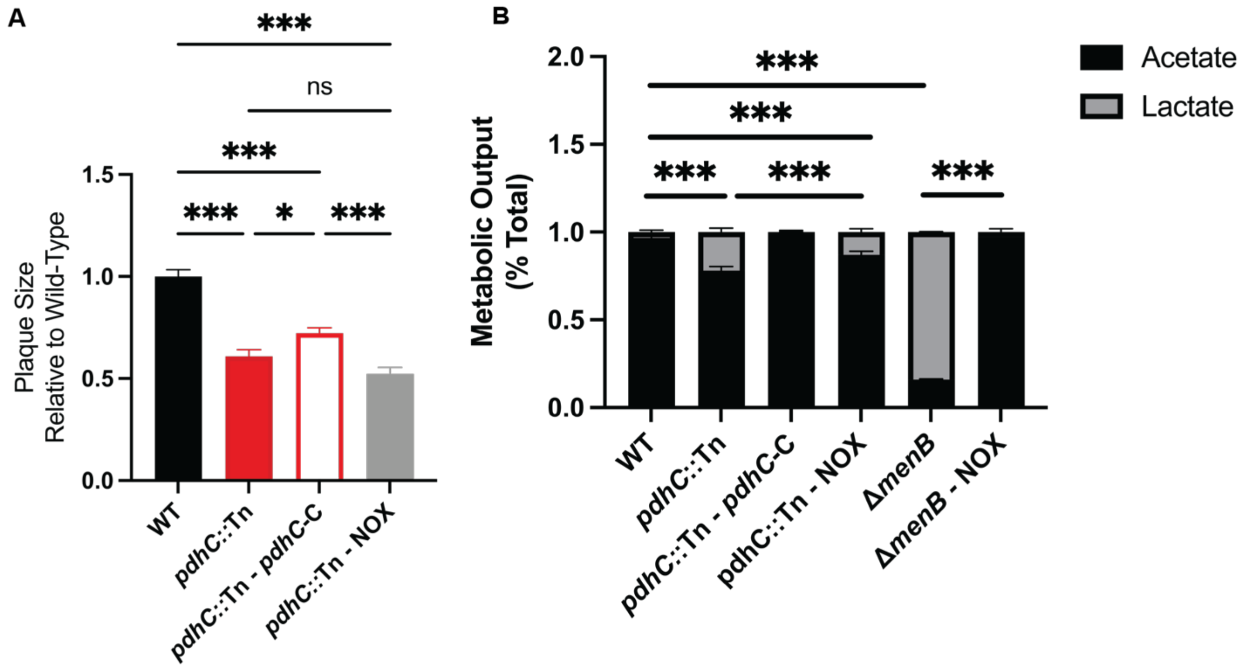


#### Supplemental Figure 1. Overexpression of NADH Oxidase, NOX, fails to rescue *pdhC*::Tn virulence as measured by plaquing assay and fermentative byproducts.

(A) L2 fibroblasts were infected with indicated *L. monocytogenes* strains at an MOI of 0.5 and were examined for plaque formation 4 days post infection. Assays were performed in biological triplicate and data displayed is the Mean and SEM of a strain's plaque size relative to WT in one of three representative biological replicates. (B) Note data for WT, *pdhC*::Tn, and *pdhC*::Tn-*pdhC*-C is recapitulation of data presented in **Figure 2**. High-performance liquid chromatography (HPLC) was used to quantify fermentation products (acetate and lactate) produced and secreted by the indicated *L. monocytogenes* strains grown aerobically in BHI medium at 37°C to stationary phase. The mean percentage of acetate and lactate production by each strain was compared to that of the wild-type *L. monocytogenes*.

#### Supplemental Table 1. Bacterial strains used in this study.

| Strain | Description | Reference |
|---|---|---|
| XL1-Blue | competent *E. coli* strain | (75) |
| S17 | *E. coli* strain for conjugations into *L. monocytogenes;* $Sp^R$ | (72) |
| 10403S [JDS 1] | Background *L. monocytogenes* 10403s strain | (77) |
| JDS 875 | *pdhA*::Tn | This Work |
| JDS 1203 | *pdhC*::Tn | (20) |
| JDS 876 | *pdhD*::Tn | This Work |
| JDS 1064 | *pdhC*::Tn – *pdhC*-C | This Work |
| MJF 363 | *pdhC*::Tn - NOX | This Work |
| MJF 368 | *pdhC*::Tn Supp 1 | This Work |
| MJF 370 | *pdhC*::Tn Supp 2 | This Work |
| MJF 372 | *pdhC*::Tn Supp 3 | This Work |
| MJF 374 | *pdhC*::Tn Supp 4 | This Work |
| MJF 376 | *pdhC*::Tn Supp 5 | This Work |
